# Co-occurrence patterns and habitat selection of the mountain hare, European hare, and European rabbit in urban areas of Sweden

**DOI:** 10.1101/2022.05.03.490451

**Authors:** Henriette Bach, Hannah Escoubet, Martin Mayer

## Abstract

Assessing the underlying mechanisms of co-occurrence patterns can be challenging as biotic and abiotic causations are hard to disentangle. To date, few studies have investigated co-occurrence patterns within urban areas that constitute novel habitat to numerous wildlife species. Moreover, as urban areas expand and are increasingly used as habitat by wildlife, there is a need for a better understanding of urban ecology to facilitate human-wildlife coexistence. Here, we investigated co-occurrence patterns and habitat selection of the European hare (*Lepus europaeus*), mountain hare (*L. timidus*), and European rabbit (*Oryctolagus cuniculus*) inside urban areas of Sweden, using joint species distribution models and generalized linear mixed models based on citizen science observations. All three species were observed within urban areas, but European hares and rabbits appear to be more successful urban colonizers compared to mountain hares. Overall, our findings suggested that urban occurrence by all three lagomorphs was related to suitable conditions within the distribution of each species (e.g. climate and elevation), rather than by the presence of other lagomorph species or specific land cover types within urban areas. On a finer spatial scale, our findings suggested facilitation of European hares by rabbits, though the mechanism for this remains unclear. European hares and rabbits generally selected for green urban areas and mountain hares for residential gardens, which likely constitute suitable foraging sites. Our findings contribute to the understanding of urban ecology and provide valuable insight for management measures of the three lagomorphs in urban areas of Sweden.

## 1. Introduction

Studying the underlying mechanisms of species co-occurrence and interactions can be challenging, because disentangling abiotic and biotic factors affecting the occurrence and abundance of species is difficult, especially in heterogeneous environments. However, studies on co-occurrence and new methods, which can untwine abiotic and biotic factors, have received more attention in recent years. Niche differences, distinct habitat preferences, competitive exclusion, environmental filtering, or a combination of these factors were proposed as ecological explanations for species segregation or co-occurrence (Pollock *et al*. 2014; Bar-Massada 2015; Estevo, Nagy-Reis & Nichols 2017; Kohli, Terry & Rowe 2018; Ulrich *et al*. 2018).

Urban areas are highly human-modified, which might lead to altered co-occurrence and interactions of the species that are able to persist in these areas. Urbanization increases globally and is a major driver of environmental change, negatively affecting ecosystems globally (Brown 2001; Grimm *et al*. 2008). Both the expansion of urban areas into animal habitats as well as active urban colonization leads to the increasing occurrence of wildlife in these novel habitats (Luniak 2004). The causes of urban colonization are often not well understood. For example, urban colonization might be driven by poorer habitat conditions or increased hunting pressure outside urban areas (Rutz 2008; Mayer & Sunde 2020). Thus, urban areas can constitute an advantageous habitat, e.g. due to relaxed predation (Møller 2012) or increased resource availability (Contesse *et al*. 2004). While some species proliferate in urban areas, others are not able to adapt and become locally extinct (McKinney 2006; Shochat *et al*. 2006). Consequently, an increased understanding of habitat preferences by urban wildlife can be a valuable tool to aid conservation actions, ensuring suitable habitats for urban colonizers. At the same time, it is important to consider biotic interactions, as competition between species might cause exclusion from otherwise suitable habitats (Thulin 2003).While habitat selection within urban areas has been previously addressed in numerous species (Chambers & Dickman 2002; Bozek, Prange & Gehrt 2007; Duduś *et al*. 2014; Mayer & Sunde 2020), research on urban community ecology and species interactions are scarce (Carrete *et al*. 2010; Magle *et al*. 2012; Ramírez-Cruz *et al*. 2019).

Citizen science observations provide large quantities of data from broad geographical scales and restricted areas (e.g. private property), which would otherwise be nigh impossible to obtain for researchers (Dickinson, Zuckerberg & Bonter 2010). However, such data also has limitations. Point occurrence data collected by volunteers is prone to pseudo-absences, temporal and spatial biases, and varying observer quality (Crall *et al*. 2011; Geldmann *et al*. 2016). Nevertheless, high human population densities within urban areas yield good coverage and increased sampling efforts (Dickinson, Zuckerberg & Bonter 2010; Mair & Ruete 2016).

Using citizen science observations, we investigated co-occurrence patterns and habitat selection of the European hare (*Lepus europaeus*), mountain hare (*L. timidus*), and European rabbit (*Oryctolagus cuniculus*, hereafter rabbit) in urban areas of Sweden to assess how occurrence and habitat selection was affected by land cover and the presence of other lagomorphs. Sweden was selected as a case study due to its high quantity of citizen science data, and for harboring three lagomorph species of similar ecology, providing an optimal model to investigate species interactions (Leach, Montgomery & Reid 2015).

Mountain hares are native to Sweden, typically associated with tundra, open forest, and heathland in upland areas (Flux & Angermann 1990; Thulin 2003). Moreover, mountain hare occurrence is positively associated with deep and lasting snow cover, and negatively with human influence (Jansson & Pehrson 2007; Leach, Montgomery & Reid 2016). They are declining and categorized as near threatened in Sweden (Artdatabanken 2020), with milder winters and competitive exclusion by European hares expanding their distribution northwards proposed to be responsible for this decline (Thulin 2003; Jansson & Pehrson 2007). Both European hares and rabbits were introduced to Sweden (Artdatabanken 2020). European hares are associated with agricultural lowland, and their densities have been found to be positively correlated with higher temperatures and lower precipitation (Smith, Jennings & Harris 2005; Leach, Montgomery & Reid 2016). While European hare populations have been declining in large parts of Europe since 1960 due to agricultural intensification (Smith, Jennings & Harris 2005), they might still be expanding their distribution in Sweden (Jansson & Pehrson 2007). The rabbit is categorized as ‘near threatened’ in its native range in the Iberian peninsula, but appears to proliferate in areas where it was introduced (Lees & Bell 2008). Although flexible in their habitat preferences, rabbits are predominantly found in grassland, pastures or arable land bordering scrubland, providing cover from predators (Calvete *et al*. 2004; Tapia *et al*. 2014). They prefer sandy soil that allows them to dig burrows, and their distribution is positively correlated with temperature and negatively with precipitation and mean slope (Calvete *et al*. 2004; Leach, Montgomery & Reid 2016). All three species are game species in Sweden, with European hares and rabbits being regulated in areas where they might cause damage (https://jagareforbundet.se/).

Previous studies are not in compliance on European hare and rabbit interactions. Most studies have found no or limited evidence for competition between the two species (Stott 2003; Katona *et al*. 2004; Flux 2008), with one study suggesting facilitation (Leach, Montgomery & Reid 2017). However, an assessment of the effectiveness of different measures to eradicate rabbits from islands showed that European hares were markedly more effective than both cats and myxomatosis in removing rabbits due to competitive exclusion (Flux 1993). The distribution of the mountain hare, apart from being affected by abiotic factors (Leach, Montgomery & Reid 2016), might be limited via competitive exclusion by the European hares’ northward expansion (Thulin 2003).

Both European hares and rabbits now occur in urban areas (Mayer & Sunde 2020; Ziege *et al*. 2020) that, under certain conditions, appear to constitute advantageous habitat. For example, rabbits became more diurnal, spent less energy on anti-predator behaviors, and reduced their home range size, possibly due to increased resource availability (Ziege *et al*. 2016; Ziege *et al*. 2020). There is little information regarding urban colonization by mountain hares, although some urban and suburban observations exist (Haigh & Lawton 2007; Levänen, Pohjoismäki & Kunnasranta 2019).

Here, we first described patterns of urban occurrence by the three lagomorphs, and then used joint distribution models to investigate the underlying mechanisms (i.e. environmental filtering or biotic interactions) of the three species’ co-occurrence patterns on urban area and 1×1 km urban grid cell level. Moreover, we investigated species occurrence and habitat selection within urban areas, assessing the role of urban area size, climate, elevation (occurrence analysis only), urban land cover types and observations of the other lagomorph species. We predicted that urbanization might increase competition for resources between European hares and rabbits, which should lead to segregation of the two species within urban areas. We further predicted that mountain hares segregate from both European hares and rabbits, due to environmental filtering, given the mountain hares’ distinct habitat preferences, and potentially due to competitive exclusion. Regarding habitat selection, we predicted that the lagomorphs selected land covers that resemble those of their preferred habitats outside urban areas, i.e. European hares and rabbits selecting open herbaceous vegetated areas, e.g. green urban areas and residential lawns (and rabbits additionally for sandy soils), and mountain hares selecting forested areas.

## 2. Methods

### Study areas and preparation of spatial data

Our study area comprised Urban Morphological Zones (UMZ) of the CORINE Land Cover 2000 version 16, defined as areas within 200 meters of each other considered to contribute to the urban tissue and function (https://www.eea.europa.eu/data-and-maps/data/urban-morphological-zones-2000-2), within Sweden, obtained from The European Environment Agency (EEA) (http://ftp.eea.europa.eu/www/umz/v4f0/UMZ2000.zip). Because higher human population densities increase sampling effort, thereby reducing the number of pseudo-absences and the effect of spatially biased sampling effort in point occurrence data (Geldmann *et al*. 2016; Mair & Ruete 2016), we only considered UMZ’s > 10 km^2^ (hereafter urban areas) for our analysis, leaving 97 urban areas (Fig. 1A). Moreover, we created 1×1 km grid cells within urban areas using ArcGISPro 2.8.3 (Esri Inc. 2020), resulting in 4,915 grid cells, to analyze species associations on a finer spatial scale (see below).

**Fig. 1:**
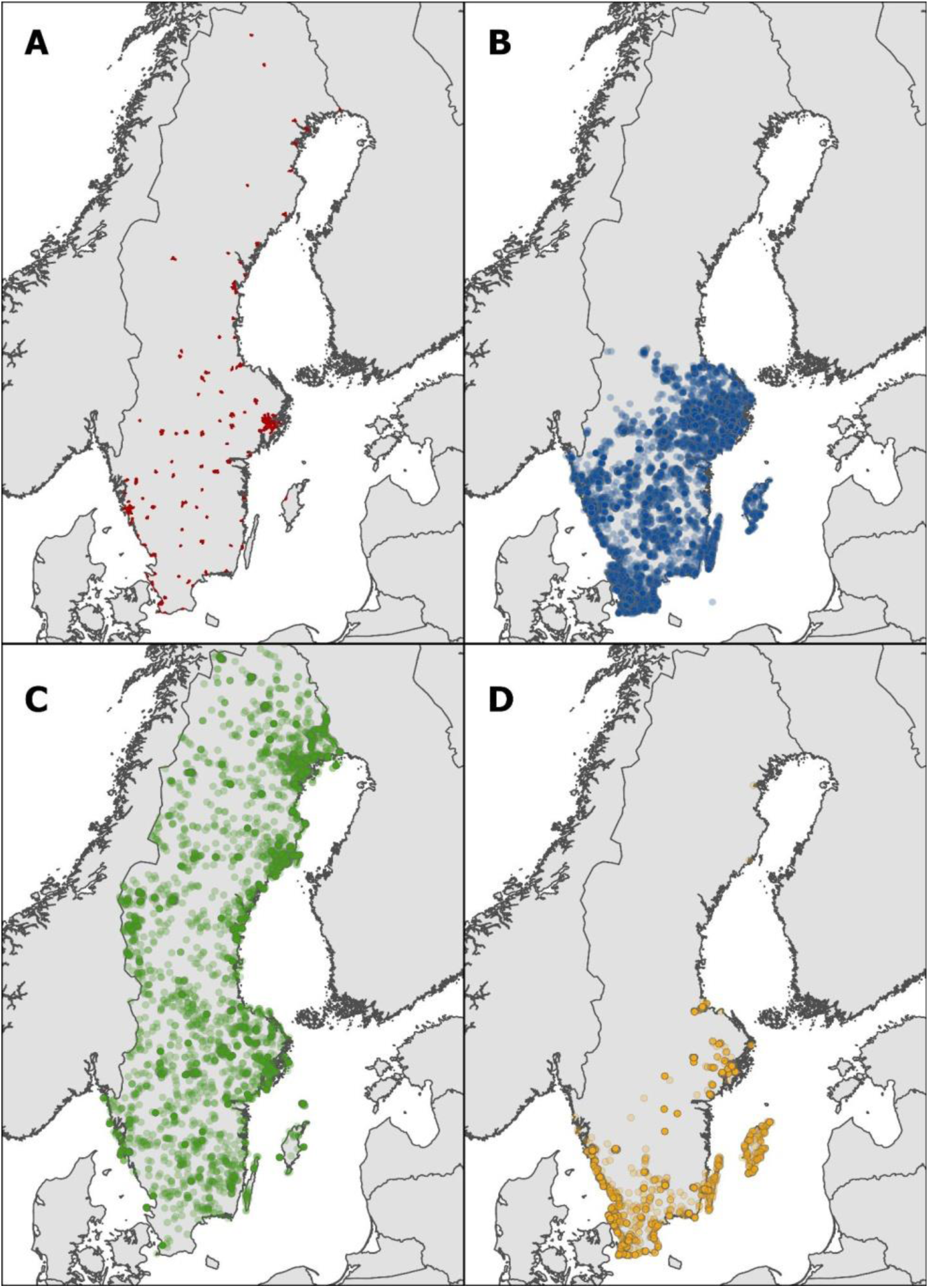
Depicting (A) the urban areas (dark red) of Sweden, and all reported citizen observations of (B) European hares (blue dots), (C) mountain hares (green dots), and (D) European rabbits (orange dots) within Sweden from 2007-2021.

### Citizen science observations and hunting bag data

Point occurrence data for the three lagomorph species within Sweden were derived from the Global Biodiversity Information Faculty (GBIF) (GBIF.org (21 March 2022) GBIF Occurrence Download https://doi.org/10.15468/dl.du6h5m) for the years 2007-2021 (Fig. 1B-D). There were very few observations before 2007, which is why we used this year as cut-off. GBIF is a database providing institutions from all over the world with common standards and open-source tools for sharing information on when and where species have been recorded (https://www.gbif.org/what-is-gbif). The bulk of data in this study (98.5%) came from Artportalen (https://www.artportalen.se/Home/About), a website for reporting species in Sweden. To reduce variation in data quality and ensure a certain degree of location precision, observations with a spatial uncertainty of >1000 m were excluded. We intersected all observations with urban areas and grid cells to assign them to environmental variables (see below), using the R package ‘raster’ (Hijmans *et al*. 2015).

Moreover, we used hunting bag data, derived from the Swedish Hunters Associations databank Viltdata (https://rapport.viltdata.se/statistik/), as a relative measure of non-urban population trends for the three species. We compared hunting bag data with the proportion of urban observations (compared to all observations) for each species separately for each year, to assess concurrence and temporal patterns in urban colonization. Changes in human urban population were obtained from the World Bank database (https://data.worldbank.org/indicator/SP.URB.TOTL.IN.ZS?locations=SE).

### Environmental data

We obtained environmental data known to and/or suspected to affect the occurrence of the three species. We obtained climate data (i.e. annual mean temperature, mean temperature of the coldest quarter, and annual precipitation) from 1970-2000 at 2.5 arc-minute resolution from WorldClim version 2.1. (https://www.worldclim.org), and mean soil sand content and bulk density at 250 meters resolution from the International Soil Reference and Information Centre version 2.0.1. (https://maps.isric.org/). We selected a depth of 15-30 cm, because rabbit burrow depth is 20 cm on average (Serrano & Hidalgo de Trucios 2011). Elevation data at 90 m resolution were obtained from the Shuttle Radar Topography Mission (http://srtm.csi.cgiar.org/). To extract climate, soil and elevation information for the urban areas, we created 100 random points per urban area using ArcGIS Pro2.8.3. We extracted the environmental values from each raster layer using the R package ‘raster’ (Hijmans *et al*. 2015) and values were averaged for each urban area. Land cover data at Minimum Mapping Unit 25 ha were downloaded from CORINE Land Cover 2018 version of the Copernicus Land Monitoring Service (https://land.copernicus.eu/pan-european/corine-land-cover/clc2018?tab=download). We intersected the land cover vector with urban areas and urban grid cells and calculated the area of each land cover patch. Moreover, to describe the land cover surrounding urban areas, we buffered each urban area by 1000 m, and then intersected this buffer with the land cover vector. We re-classified the CORINE land cover classes into 8 categories within urban areas: (1) continuous urban fabric (e.g. city centers, >80% of the ground covered by artificial surfaces, i.e. soil sealed), (2) discontinuous urban fabric (e.g. suburbs, >30% scattered urban fabric without sealed soil), (3) industry (including airports, railways, etc.), (4) green urban areas (e.g. parks), (5) agriculture, (6) forest (including other (semi)natural areas including heathland), (7) water, and (8) other areas (water, beaches, bare rock, etc.) (Table S1). The land covers surrounding urban areas were categorized into (1) agriculture, (2) forest, (3) urban areas (merging the above-mentioned urban categories due to little urban land cover surrounding urban areas), and (4) other areas. We then calculated the proportion of each land cover category per urban area, grid cell, and surrounding urban areas.

### Defining species occurrence

We categorized a species occurring in an urban area when there were ≥7 observations within the urban area, i.e. at least one observation per year when most observations were recorded (from 2015-2021; see results). We chose this categorization to minimize defining species occurrence based on misidentifications, observations of escaped/released pet hares and rabbits, or dispersing individuals that had not established in the area. Defining occurrence based on a single observation did not markedly change the results (not shown). On grid cell level, we defined species presence in grid cells with ≥1 observation, because on this fine scale, point occurrence data likely was more prone to false-negatives rather than false-positives (Crall *et al*. 2011).

### Data analyses

First, to assess whether there was evidence of biotic interactions between the three species and whether the three species shared or had distinct environmental affiliations, we used the joint species distribution model (JSDM) provided by Pollock *et al*. (2014), which accounts for co-occurrence patterns of multiple species. We modeled predicted probabilities of occurrences, using a binary response variable (i.e. presence/absence), using a multivariate probit regression model (Pollock *et al*. 2014). The model provides environmental correlations for species pairs indicating shared or differing environmental responses, whilst residual correlations suggest biotic interactions, such as competition or facilitation (Pollock et al. 2014). We did two model runs, one on an urban area level and one on a grid cell level, as environmental effects and competitive interactions are known to appear at different scales (Leach, Montgomery & Reid 2017). For the JSDMs, collinearity among environmental variables was assessed building a correlation matrix, defining correlation as Pearson’s coefficient > 0.6 (Zuur, Ieno & Elphick 2010). Consequently, we removed mean annual temperature (positively correlated with temperature of coldest quarter), soil bulk density (positively correlated with soil sand content) and surrounding forest (negatively correlated with surrounding agriculture). Moreover, we removed the proportion of forest (positively correlated with surrounding forest and negatively with surrounding agriculture) for the urban area level analysis and the discontinuous urban fabric (negatively correlated with industry) for the analysis on grid cell level. Consequently, we included temperature of coldest quarter, soil sand content, precipitation, elevation, the proportion of continuous urban fabric, discontinuous urban fabric (for the urban area level analysis only), forest (for the grid cell level analysis only), industry, green urban areas, agriculture, surrounding agriculture and surrounding urban fabric. We centered and scaled covariates prior to analyses (Grueber *et al*. 2011).

Second, we investigated the factors affecting species occurrence within urban areas, separately for the three lagomorphs within their distribution, using generalized linear models with a log link and a binomial response distribution (present = 1 versus absent = 0). To estimate the European hares’ and rabbits’ distribution in Sweden (the mountain hares’ range covers the whole of Sweden), we created 100% minimum convex polygons based on citizen science observations (excluding obvious outliers). We initially built 4 candidate models based on biological hypotheses: (1) land covers within urban areas affect the presence of a species, including the proportion of continuous urban fabric, discontinuous urban fabric, green urban areas, industry, forest, and agriculture; (2) land covers surrounding urban areas affect species occurrence, including the proportion of surrounding agriculture, forest, and urban areas; (3) climate and the size of an urban area (as proxy for the number of observers) affect occurrence, including mean temperature of coldest quarter, mean annual precipitation, elevation, soil sand content (for rabbits only), and urban area size; (4) competition with or facilitation by other lagomorphs affects occurrence, including the presence of other lagomorph species (estimated as above). The proportion of surrounding forest and agriculture were highly correlated (Pearson’s correlation coefficient > 0.6 and variance inflation factor > 3 (Zuur, Ieno & Elphick 2010)) so we only included agriculture (European hares and rabbits) or forest (mountain hares) in our analysis. Additionally, as measure of relative abundance, we analyzed the number of observations per urban area separately for each species (again including all urban areas within the species’ distribution), using generalized linear models of the R package ‘glmmTMB’ (Magnusson *et al*. 2017) with a log link function and negative binomial distribution to account for overdispersion and zero-inflation (O’hara & Kotze 2010). We again built the same candidate models as for the species presence analyses. We scaled all numeric variables (mean = 0; standard deviation = 1) to obtain comparable estimates. We initially compared the 4 models based on biological hypotheses for both the analyses of species presence and relative abundance using Akaike’s Information Criterion (AIC). To obtain the most parsimonious (hereafter best) model, we performed a stepwise backward selection, starting from the full model including all variables, and removed variables that lead to an increase in AIC, selecting the model with the lowest AIC (Wagenmakers & Farrell 2004). This approach resulted in the same best model compared to model selection based on creating all possible combinations of candidate models using the ‘dredge’ function of the R package ‘MuMIn’ (Barton 2020).

Third, to analyze habitat selection within urban areas, we selected urban areas that had at least 10 observations of a given species. For this analysis, we excluded observations with spatial uncertainty of >500 m, because this analysis was conducted at a finer spatial scale. To get a measure of resource availability, we created 5× the number of random positions than we had obtained from citizen observations within each urban area. We then assigned each random and used (observed) position to the land cover type (as defined above) and the soil sand content (for rabbits only). To analyze habitat selection (observed location = 1 versus random location = 0, dependent variable), we used generalized linear mixed models with a binomial distribution and a logit link, using the R package ‘lme4’ (Bates *et al*. 2015). We included the land cover type (excluding ‘other’ land cover), soil sand content (for rabbits only), the presence of other lagomorph species, and the interaction of land cover type with lagomorph presence (to investigate if habitat selection differs in the presence of other lagomorphs) as fixed effects, and urban ID as random intercept to control for non-independence of the data. We again performed a stepwise backward selection, starting from the full model including all variables, selecting the model with the lowest AIC. Parameters that included zero within their 95% confidence interval were considered uninformative (Arnold 2010). All analyses were carried out in R4.0.3.

## 3. Results

### Patterns of urban occurrence

Out of a total of 20,931 observations (both within and outside urban areas), European hares constituted 12,492 observations (60%), mountain hares 5,727 observations (27%), and rabbits 2,712 observations (13%). The number of observations increased over time, with few observations before 2015, when Artportalen was initiated (Fig. 2A). Of the three species, rabbits had the highest proportion of urban observations, accounting for 39% (1,049 observations) of all rabbit observations. The proportion of urban rabbit observations fluctuated between years, with noticeable decreases in 2008-2009, 2014-2015 and 2020 (Fig. 2B). For European hares, 22% (2,769) were urban observations, with the proportion of urban observations being relatively stable over time (Fig. 2B). For mountain hares, urban observations accounted for 12% (714 observations), and the proportion of urban observations increased over the years, being 2.4% in 2007 and 21% in 2021 (Fig. 2B). Hunting bag numbers decreased for European hares and mountain hares, and fluctuated for rabbits, with pronounced increases in 2009 and 2015 (Fig. 2C). The percentage of people living in urban areas increased from 85% in 2007 to 88% in 2020.

**Fig. 2:**
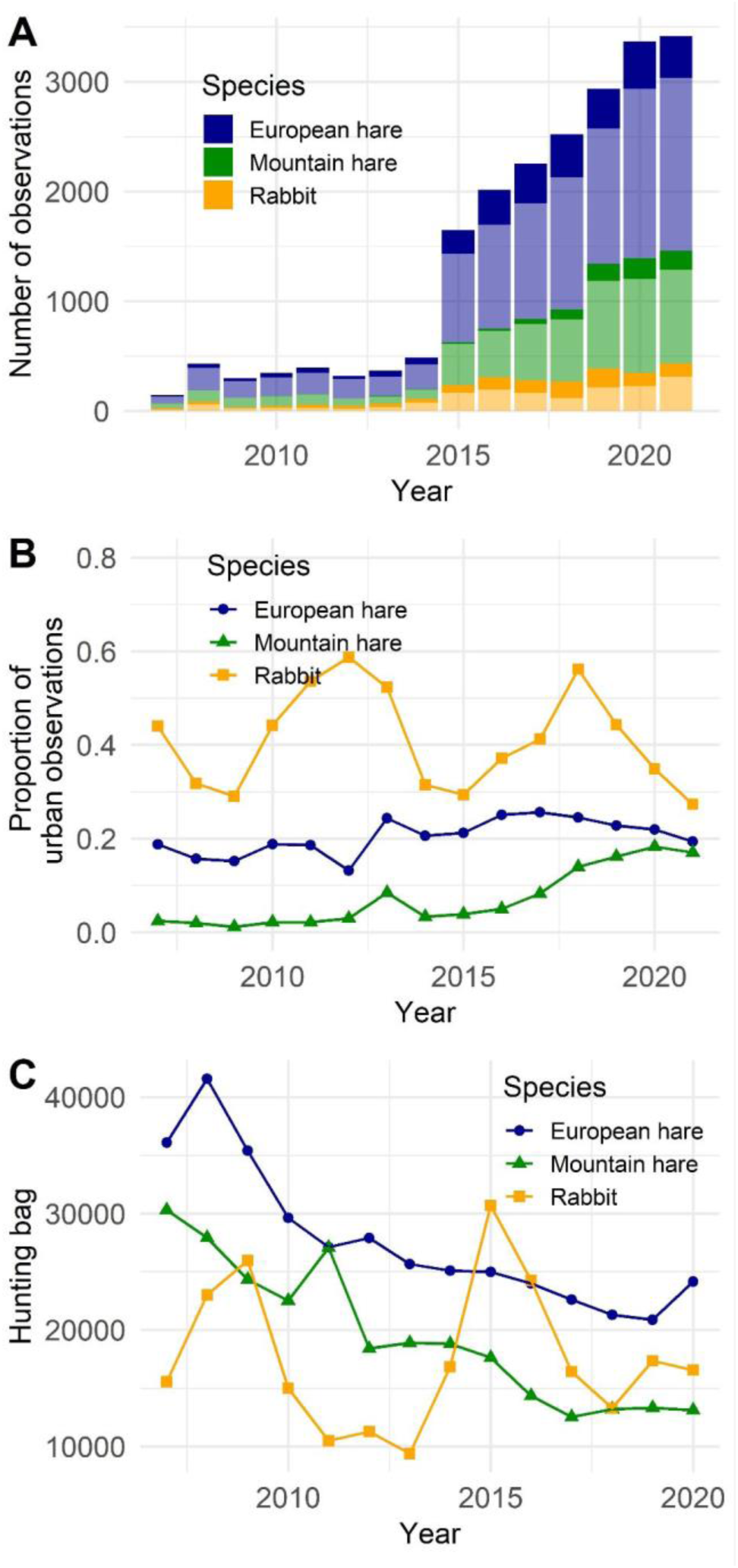
(A) The number of all reported observations for each year (2007-2021) and species, and shown for urban observations (dark colors) and observations outside urban areas (bright colors). (B) The proportion of urban observations out of total observations for the three species, and (C) hunting bag numbers from 2007 to 2020 (numbers from 2021 were not available yet).

### Environmental and residual correlations

European hares and rabbits shared environmental responses (i.e. concerning mean temperature of the coldest quarter, annual precipitation, soil sand content, elevation, and land cover proportions), while mountain hares had distinct environmental responses from the other two species (Table 1, Fig. 3). These responses were more pronounced on a grid cell level. All three species pairs had positive residual correlations (especially European and mountain hares, and rabbits and mountain hares), suggesting that all species pairs co-occurred more than expected, due to unmodelled factors (Table 1, Fig. 3). For European and mountain hares, and rabbits and mountain hares, this pattern was more pronounced on urban level compared to grid cell level, but for European hares and rabbits residual correlation was stronger on grid cell level (Table 1).

**Fig. 3:**
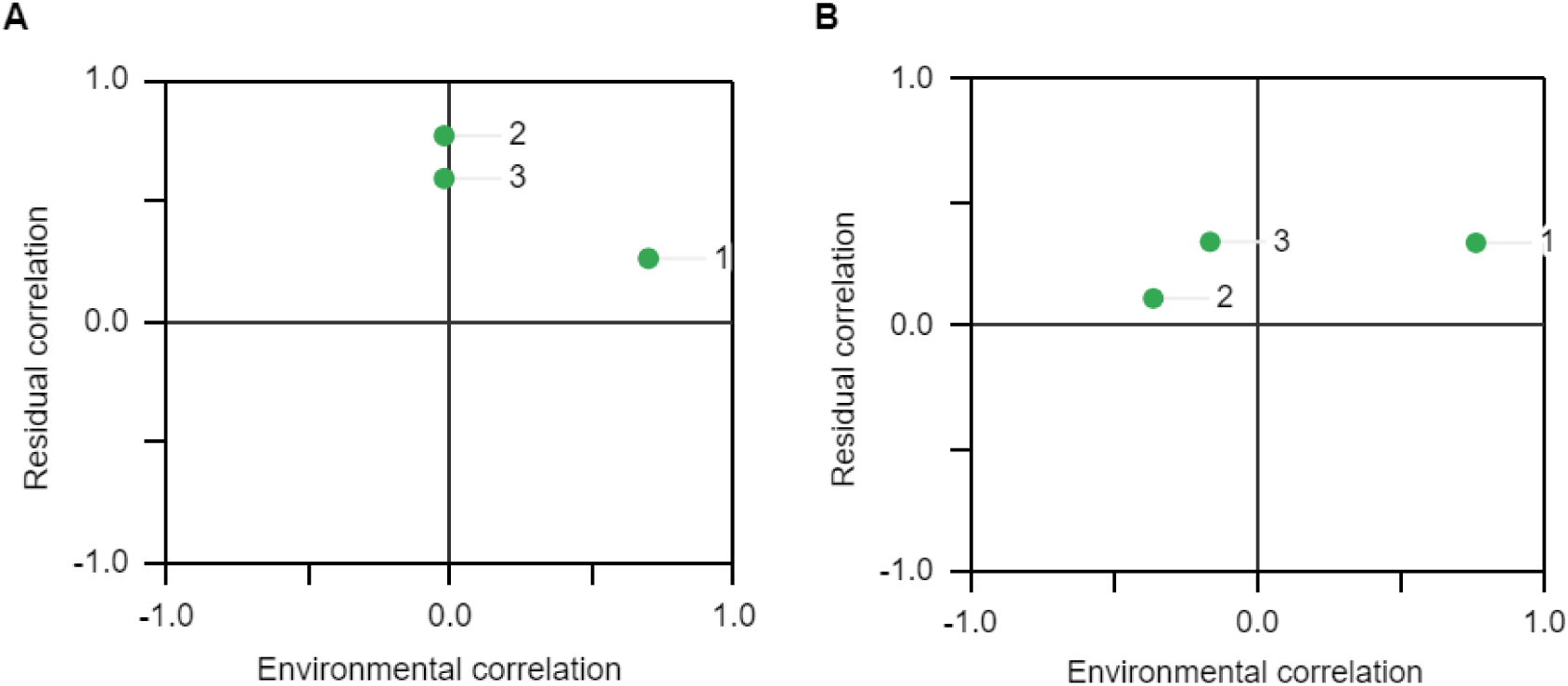
Environmental and residual correlations between European hare and European rabbit (1), European rabbit and mountain hare (2), and mountain hare and European hare (3), on urban area level (A), and grid cell level (B).

**Table 1:**
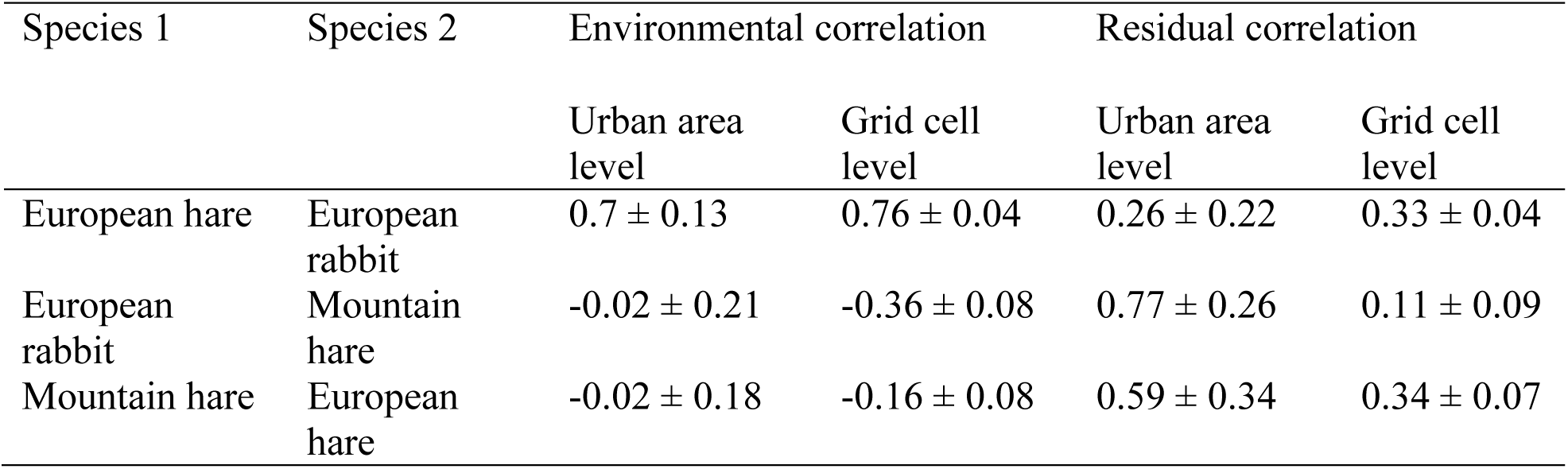
Mean (± SD) environmental and residual correlations between the three species pairs separately for urban and grid cell level.

### Species occurrence and observations per urban area

Within their respective ranges (Fig. 1), 63 of 77 urban areas (82%) contained European hare observations, 38 of 97 urban areas (39%) contained mountain hare observations, and 40 of 69 urban areas (58%) contained rabbit observations. When defining species presence as at least 7 observations within an urban area, European hares occurred in 45% of urban areas within their distribution, mountain hares in 10%, and rabbits in 26% of urban areas.

For all three species, the probability of occurrence within an urban area was best explained by the model including climate variables, elevation, and the size of the urban area, followed by the model including the surrounding land cover (European and mountain hare analysis) or other species (rabbit analysis), and finally the model including land cover within urban areas (Table 2). After model selection, the best model explaining the probability of urban European hare occurrence included urban area size (positive correlation), elevation (positive correlation; Fig. 4A), mean annual precipitation (negative correlation; Fig. 4B), and temperature of the coldest quarter (uninformative positive correlation; Table 3). The probability of urban mountain hare occurrence also increased with urban area size, and declined with increased temperature of the coldest quarter (Fig. 4C, Table 3). Elevation was included in the best model, but was uninformative (positive correlation). The probability of urban rabbit occurrence also increased with urban area size, and with the proportion of green urban areas (Fig. 4D), though this effect was uninformative (Table 3).

**Fig. 4:**
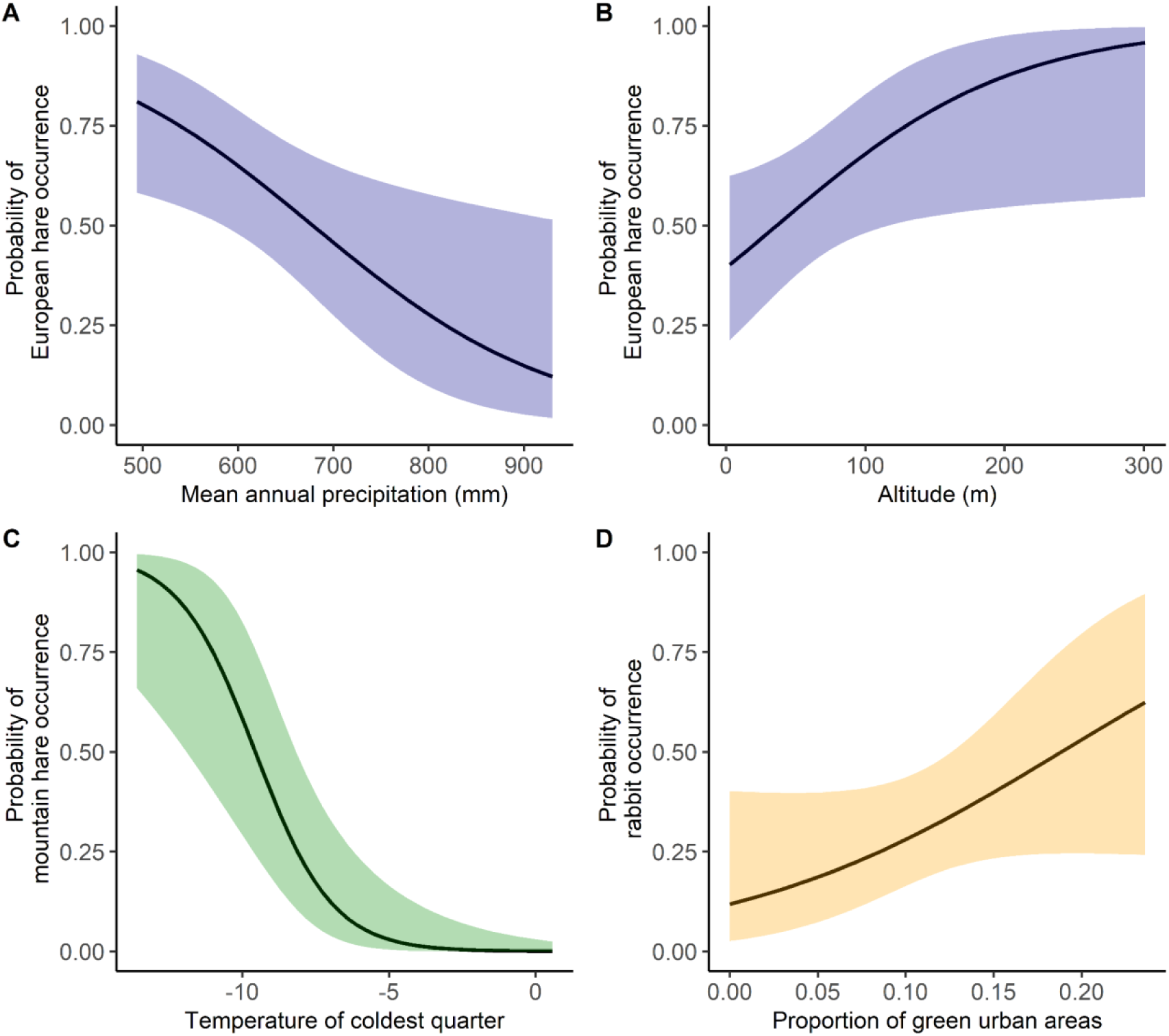
The predicted probability of urban occurrence by (A) European hares in relation to mean annual precipitation and (B) elevation, and (C) by mountain hares in relation to the temperature of the coldest quarter, and (D) by rabbits in relation to green urban areas. 95% confidence intervals are shown as shading.

**Table 2:**
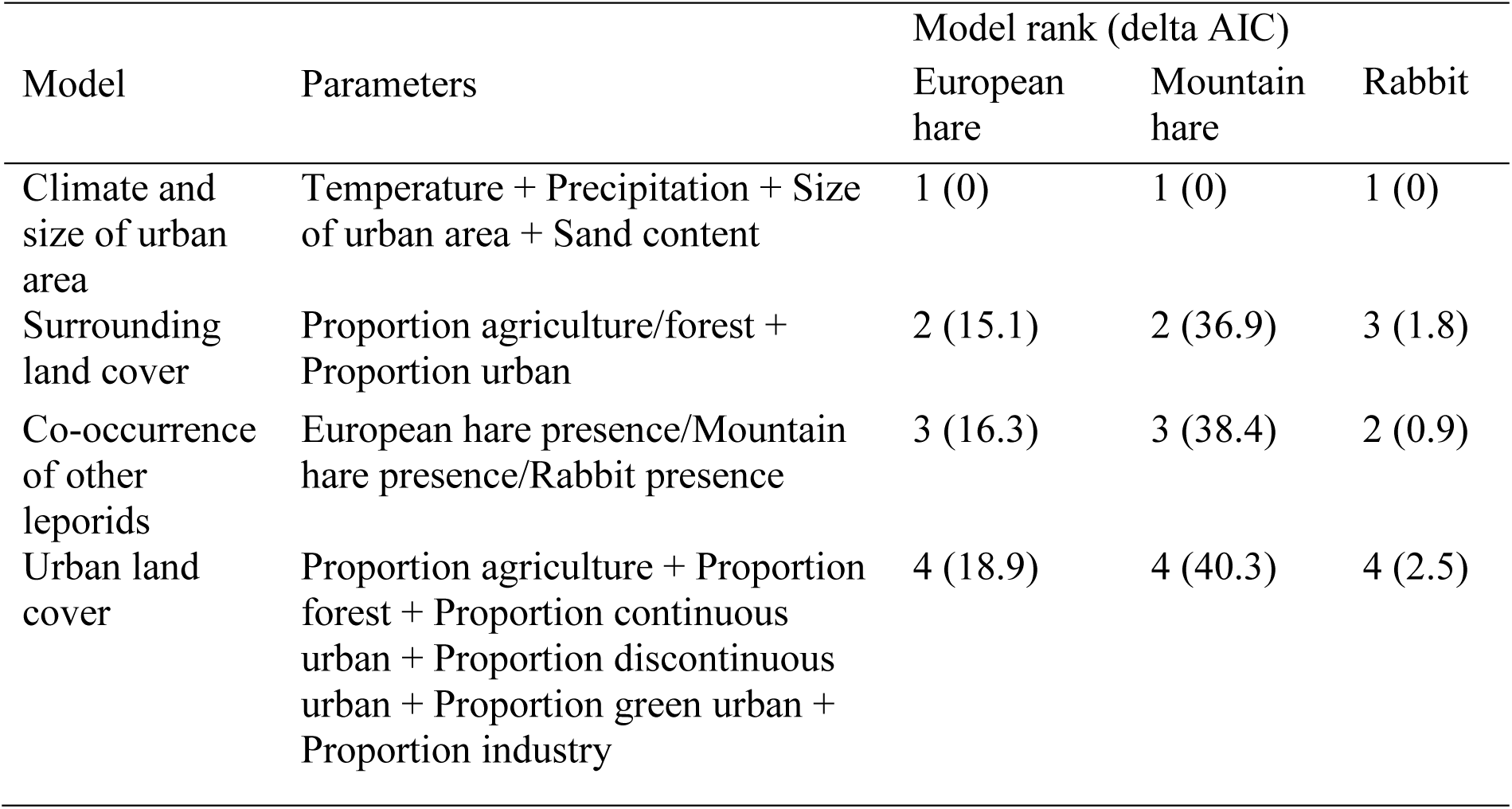
Overview of the candidate models based on biological hypothesis for the analysis of urban occurrence by European hares, mountain hares, and European rabbits. Models were ranked based on AIC.

**Table 3:** Estimate, standard error (SE), lower 95% confidence interval (LCI) and upper 95% confidence interval (UCI) of explanatory variables for the analyses of urban occurrence separately for European hares, mountain hares, and European rabbits. Informative parameters are in bold.

| Parameter | Estimate | SE | LCI | UCI |
| --- | --- | --- | --- | --- |
| *European hare occurrence* |  |  |  |  |
| Intercept | 0.31 | 0.34 | -0.34 | 1.03 |
| **Mean annual precipitation** | **-0.75** | **0.32** | **-1.42** | **-0.17** |
| **Size of the urban area** | **4.06** | **1.33** | **1.68** | **6.94** |
| **Altitude** | **0.74** | **0.36** | **0.07** | **1.51** |
| Temperature of the coldest quarter | 0.58 | 0.35 | -0.08 | 1.30 |
| *Mountain hare occurrence* |  |  |  |  |
| Intercept | -4.97 | 1.45 | -9.05 | -2.90 |
| **Temperature of the coldest quarter** | **-2.78** | **0.92** | **-5.21** | **-1.37** |
| **Size of the urban area** | **4.19** | **2.86** | **1.44** | **11.24** |
| Altitude | 0.95 | 0.61 | -0.04 | 2.65 |
| *Rabbit occurrence* |  |  |  |  |
| Intercept | -0.93 | 0.35 | -1.65 | -0.26 |
| Proportion of green urban areas | 0.57 | 0.35 | -0.08 | 1.30 |
| **Size of the urban area** | **2.71** | **1.19** | **0.68** | **5.23** |

The number of observations per urban area ranged from 0 to 846 (mean ± SD = 36 ± 116, median = 5) for European hares, from 0 to 461 (mean ± SD; 8 ± 48, median = 0) for mountain hares, and from 0 to 226 (mean ± SD; 15 ± 40, median = 1) for rabbits. The number of European hare observations per urban area was positively correlated with the size of the urban area, the proportion of forest, continuous urban fabric, surrounding agriculture, and rabbit presence, and negatively correlated with increasing precipitation and temperature of the coldest quarter (Table S2, Table S3, Fig. S1). Urban mountain hare observations were positively correlated with urban area size, the proportion of surrounding urban areas, and European hare presence, and negatively with the proportion of discontinuous urban fabric and temperature of the coldest quarter (Table S2, Table S3, Fig. S2). Proportion of agriculture was included in the best model (positive correlation), but was uninformative. Urban rabbit observations were positively correlated with increasing urban area size, soil sand content, proportion of discontinuous urban fabric, green urban areas, industry, and the proportion of surrounding urban areas, and negatively with increasing elevation and proportion of urban forest (Table S2, Table S3, Fig. S3).

### Habitat use and selection within urban areas

Based on random positions (located within urban areas where lagomorphs were present), urban areas were dominated by discontinuous urban fabric (61%), followed by industrial areas (20%), green urban areas (13%), forest (3%), continuous urban fabric (<2%), and agriculture (<2%). All three species were mostly observed in discontinuous urban fabric (especially mountain hares), followed by green urban and industrial areas (Fig. 5A). For any species, <5% of observations came from continuous urban fabric, forest, and agriculture combined. Further, there were more European hare and rabbit observations in discontinuous urban fabric when they occurred in absence of the other lagomorph species (Fig. 5A). The opposite was the case in green urban and industrial areas, i.e. more European hares were observed in green urban areas when rabbits were present, and more rabbits were observed in industrial areas when European hares were present (Fig. 5A).

**Fig. 5:**
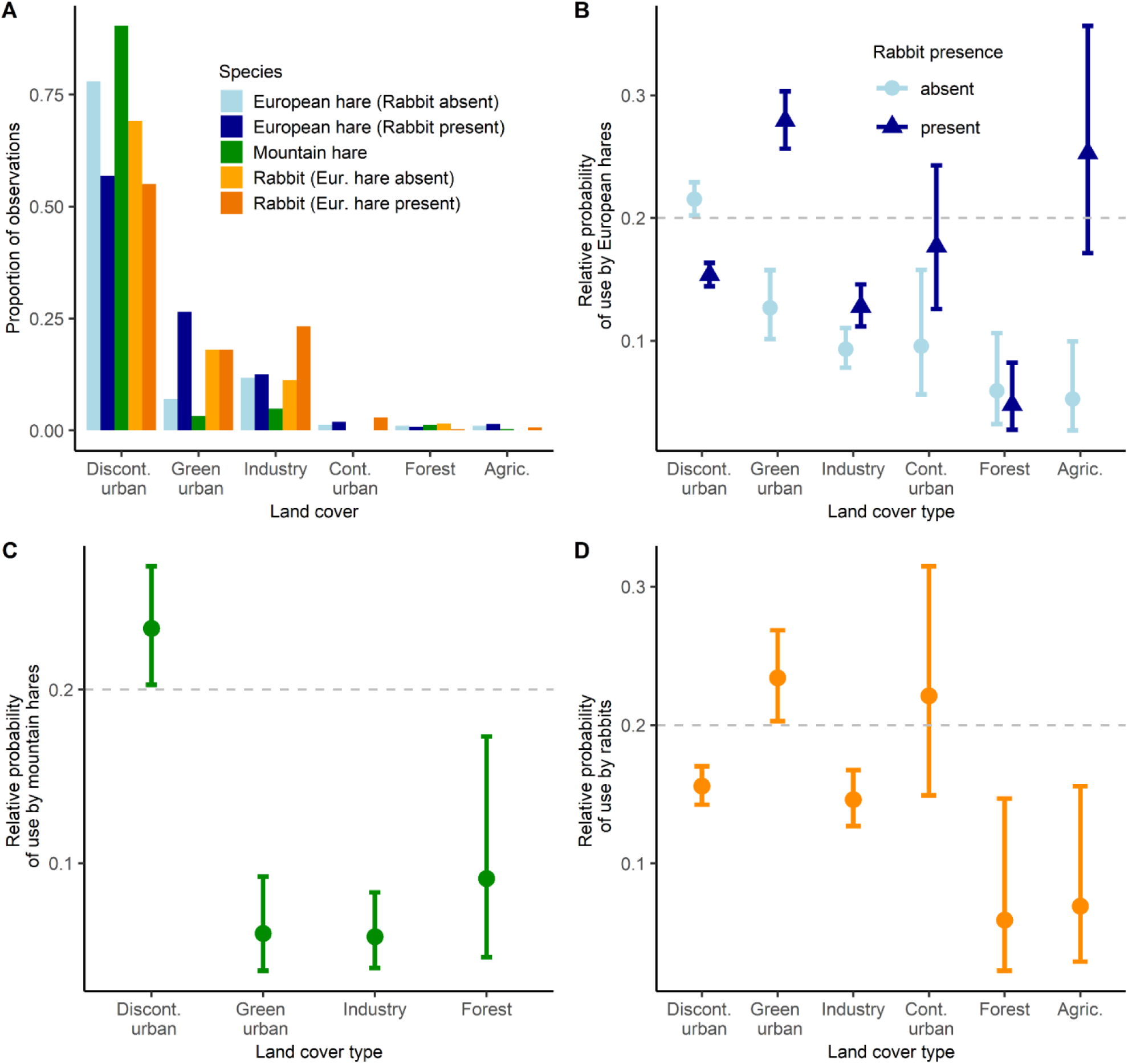
(A) The proportion of urban observations in the different land cover categories separately for the three species. For European hares and rabbits, observations are further separated by the presence or absence of rabbits/European hares (mountain hares were only observed in urban areas without the other two species). Moreover, the relative probability of use by European hares (B), mountain hares (C), and rabbits (D). For European hares, rabbit presence affected habitat selection, but not for mountain hares and rabbits. Values >0.2 indicate selection, whereas values <0.2 indicate avoidance. The 95% confidence intervals are given as bars.

Habitat selection by European hares differed between urban areas where rabbits were absent versus present (Table 4). European hares selected for green urban areas and avoided discontinuous urban fabric when rabbits were present, but avoided green areas and selected for discontinuous urban fabric when rabbits were absent (Fig. 5B). Moreover, they showed no clear selection or avoidance of continuous urban fabric and agriculture when rabbits were present, but avoided these land covers when rabbits were absent (Table 4). They consistently avoided industrial areas and forests independent of rabbit presence (Fig. 5B). Only 6 urban areas had at least 10 mountain hare observations, all located outside the distribution of European hares and rabbits. Mountain hares selected for discontinuous urban fabric, and avoided green urban and industrial areas, and forests (Fig. 5C, Table 4). We removed the continuous urban fabric and agriculture from this analysis, because there were no mountain hare observations in these areas and they constituted a negligible portion of the area (<1%). Habitat selection by rabbits was not affected by European hare presence. Rabbits selected for green urban areas, showed no clear selection or avoidance of continuous urban fabric, and avoided discontinuous urban fabric, industrial areas, forests and agriculture within urban areas (Fig. 5D, Table 4).

**Table 4:**
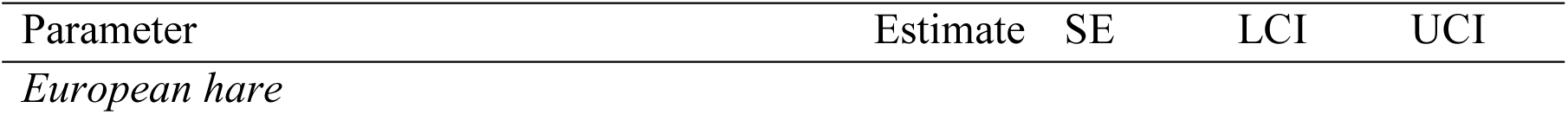
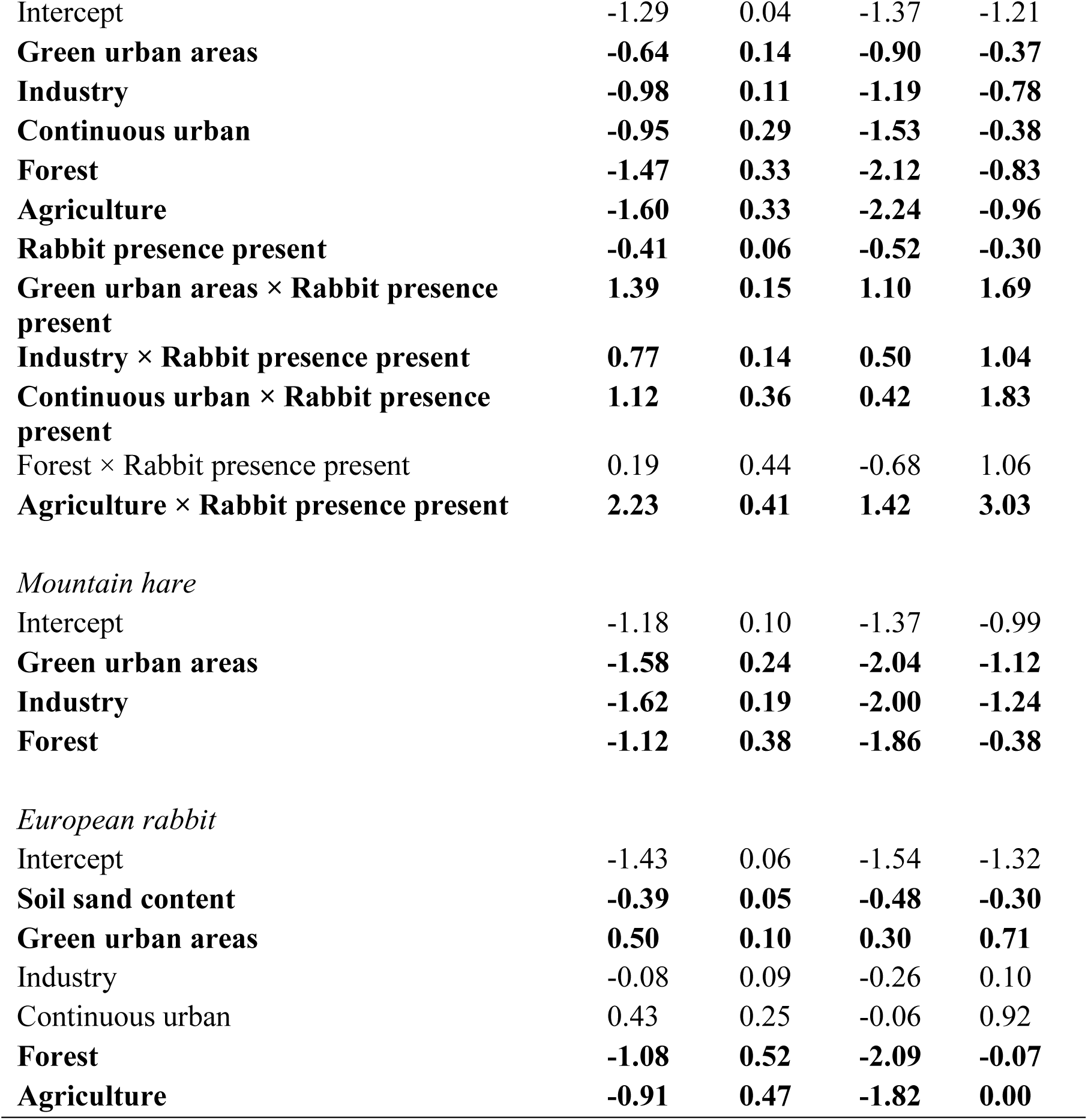
Estimate, standard error (SE), lower 95% confidence interval (LCI) and upper 95% confidence interval (UCI) of explanatory variables for the analyses of urban habitat selection separately for European hares, mountain hares and rabbits. Informative parameters are in bold. The land cover ‘discontinuous urban fabric’ was used as reference category, with positive estimates indicating a higher relative probability of use (selection) and negative values indicating a lower relative probability of use (avoidance) in comparison to this land cover.

| Parameter | Estimate | SE | LCI | UCI |
| --- | --- | --- | --- | --- |
| *European hare* |  |  |  |  |
| Intercept | -1.29 | 0.04 | -1.37 | -1.21 |
| **Green urban areas** | **-0.64** | **0.14** | **-0.90** | **-0.37** |
| **Industry** | **-0.98** | **0.11** | **-1.19** | **-0.78** |
| **Continuous urban** | **-0.95** | **0.29** | **-1.53** | **-0.38** |
| **Forest** | **-1.47** | **0.33** | **-2.12** | **-0.83** |
| **Agriculture** | **-1.60** | **0.33** | **-2.24** | **-0.96** |
| **Rabbit presence present** | **-0.41** | **0.06** | **-0.52** | **-0.30** |
| **Green urban areas × Rabbit presence present** | **1.39** | **0.15** | **1.10** | **1.69** |
| **Industry × Rabbit presence present** | **0.77** | **0.14** | **0.50** | **1.04** |
| **Continuous urban × Rabbit presence present** | **1.12** | **0.36** | **0.42** | **1.83** |
| Forest × Rabbit presence present | 0.19 | 0.44 | -0.68 | 1.06 |
| **Agriculture × Rabbit presence present** | **2.23** | **0.41** | **1.42** | **3.03** |
| *Mountain hare* |  |  |  |  |
| Intercept | -1.18 | 0.10 | -1.37 | -0.99 |
| **Green urban areas** | **-1.58** | **0.24** | **-2.04** | **-1.12** |
| **Industry** | **-1.62** | **0.19** | **-2.00** | **-1.24** |
| **Forest** | **-1.12** | **0.38** | **-1.86** | **-0.38** |
| *European rabbit* |  |  |  |  |
| Intercept | -1.43 | 0.06 | -1.54 | -1.32 |
| **Soil sand content** | **-0.39** | **0.05** | **-0.48** | **-0.30** |
| **Green urban areas** | **0.50** | **0.10** | **0.30** | **0.71** |
| Industry | -0.08 | 0.09 | -0.26 | 0.10 |
| Continuous urban | 0.43 | 0.25 | -0.06 | 0.92 |
| **Forest** | **-1.08** | **0.52** | **-2.09** | **-0.07** |
| **Agriculture** | **-0.91** | **0.47** | **-1.82** | **0.00** |

## 4. Discussion

Citizen observations were useful in describing urban occurrence and habitat selection of the three lagomorphs in Sweden. The data suggests that European hares and rabbits are successful urban colonizers, and mountain hares also begin to establish populations in some urban areas in the northern part of Sweden. Urban occurrence by all species was generally better explained by climatic conditions, elevation, and urban area size, rather than by the proportion of land cover types within urban areas or the presence of other lagomorph species. Thus, urban colonization was likely driven by suitable conditions within the distribution of each species. In contrast to our prediction, the JSDM and habitat selection analyses indicated no direct competition among the three species, but actually indicated a facilitative relationship between European hares and rabbits.

### Trends in urban observations

Both citizen observations and hunting bag data suggest that European hares were the most abundant of the three lagomorphs in Sweden. However, hunting bag reports indicated that European hare populations are declining, a trend seen throughout Europe (Smith, Jennings & Harris 2005). The relatively stable proportion of urban European hare observations over time suggests that European hare populations have established in urban areas of Sweden, similar to urban areas in Denmark (Mayer & Sunde 2020). Like European hares, rabbits appeared to be strong urban colonizers, with nearly 40% of all observations coming from urban areas, consistent with previous findings showing that rabbits are successful urban colonizers (Ziege *et al*. 2015; Ziege *et al*. 2016; Ziege *et al*. 2020). Assuming hunting bag data to be a measure of population trends, rabbit populations fluctuated over the years. Hunting bag reports and the proportion of urban observations mirrored each other well, i.e. increases in hunting bag were accompanied by decreases in the proportion of urban observations. A potential explanation might be that rabbits were culled in urban areas to prevent damage to city parks. For example, in Stockholm 6,000 rabbits were culled in 2008 and 3,000 in 2009 (https://abcnews.go.com/International/rabbits-burned-fuel-sweden/story?id=8824540), which coincided with the decrease in proportion of urban rabbit observations. Additionally, fluctuations in rabbit numbers could be related to fluctuations in climate conditions and/or disease outbreaks (Calvete *et al*. 2002; Rödel & Dekker 2012). Based on hunting bag data, mountain hare numbers were intermediate compared to European hares and rabbits, and were declining, consistent with the species’ red list status in Sweden (Artdatabanken 2020). This decline has been attributed to climate warming and competitive exclusion by and hybridization with the European hare (Thulin 2003). The proportion of urban mountain hare observations increased in recent years, indicating that urban areas are increasingly colonized by mountain hares. However, this increase might also be partly related to an increased proportion of humans living in urban areas. Overall, urban mountain hare observations were less common compared to the other two species, consistent with findings showing that mountain hares select for areas of low human influence (Leach, Montgomery & Reid 2016). Conversely, the proportion of urban observations for both European hares and rabbits might be biased in relation to mountain hare observations, because their range covered the more densely populated south of the country, potentially leading to a comparatively greater sampling effort inside urban areas (Geldmann *et al*. 2016).

### Biotic interactions and environmental filtering

European hares and rabbits shared environmental responses, while mountain hares had distinct environmental responses, consistent with previous findings (Leach, Montgomery & Reid 2017). This was likely related to the distribution of the three species, with the European hares and rabbits’ southern distribution characterized by higher temperatures and lower elevations compared to northern Sweden, where only mountain hares occurred. Both European hares and rabbits are generally associated with comparatively warm and dry climate, and lowland areas (Calvete *et al*. 2004; Smith, Jennings & Harris 2005; Tapia *et al*. 2014; Leach, Montgomery & Reid 2016), whereas mountain hares typically occupy colder areas at higher elevations (Thulin 2003; Jansson & Pehrson 2007). For all species pairs, environmental correlations were stronger on a grid cell level, probably because this finer spatial scale captured more detailed environmental differences.

The positive residual correlations (both on urban area and grid cell level) between European hares and rabbits suggest that the two species co-occurred more than expected from their shared environmental responses, indicating a facilitative interaction consistent with previous studies (Leach, Montgomery & Reid 2017). Although there is evidence that European hares and rabbits are not in competition (Stott 2003; Katona *et al*. 2004), the study by Leach, Montgomery and Reid (2017) and this study, to our knowledge, are the only implying a facilitative interaction between European hares and rabbits. Co-existence between the two species have been proposed to be mediated by the larger home range of the European hare, which enables local scale avoidance, and diet partitioning with regards to grass species (Stott 2003; Lush, Ward & Wheeler 2017). Alternatively, positive residual correlations, representing unmodelled correlations, could also represent shared environmental preferences from environmental variables not included in the models or biases in citizen observations (also see discussion of habitat selection below). Observation biases were likely, considering the high residual correlation between mountain hares and the other two species, despite the fact that the 6 urban areas with >10 mountain hare observations were all located in areas outside the other species distribution.

### Species occurrence, relative abundance and habitat selection

Urban area size was the most important factor explaining the occurrence of all three lagomorphs. This might indicate that urban areas have to be large enough to allow a sufficient number of individuals to adjust (either via selection of bold individuals or behavioral adaptations) to the novel conditions (e.g. high level of human disturbance), and consequently establish a population. Alternatively, there might not be sufficient observers in smaller urban areas to reliably detect the presence of a species, cautioning against interpreting this finding too much in the absence of a true measure of observation effort (Kelling *et al*. 2015). Apart from urban area size, the probability of urban European hare occurrence decreased with higher precipitation and tended to increase with higher temperatures, suggesting that warmer and drier areas generally favor European hare occurrence (Smith, Jennings & Harris 2005; Leach, Montgomery & Reid 2016). Moreover, the probability of European hare occurrence increased with elevation; a counterintuitive finding, as this species is typically associated with lowland. However, the average elevation of urban areas within the European hares’ distribution was 62 m, and only a single urban area was located >210 m asl (at 300 m), i.e. all urban areas were located at comparatively low elevations. The probability of mountain hare occurrence markedly decreased when temperatures were higher, in line with this species’ preference for colder climates (Jansson & Pehrson 2007). Rabbit occurrence, apart from urban area size, tended to increase when more green urban areas were present, suggesting that parks and other green areas constitute important habitat for this species. The general lack of urban land cover in the best models explaining the probability of urban occurrence suggests that factors explaining the general distribution of the species (climate and elevation) are better at predicting urban occurrence, especially for European and mountain hares. We have no evidence that species competition affected urban occurrence by any of the three species.

The analyses of the number of citizen observations per urban area yielded different results compared to the urban occurrence and habitat selection analyses. For example, the number of mountain hare observations decreased with the proportion of discontinuous urban fabric, whereas the habitat selection analysis indicated that mountain hares selected for this land cover type. Similar contrasting results were found for European hares in relation to forest and for rabbits concerning discontinuous urban fabric. We deem the analyses of relative abundance less reliable, because the number of observations was likely more biased (based on observer distribution) compared to a presence/absence measure and compared to accounting for availability in the habitat selection analysis, though the latter might have also resulted in biases due to creating random positions in areas where no observers went. This highlights that using different analytical approaches can be useful to test the generality of findings, especially when using heterogeneous citizen science data.

Inside urban areas, European hares selected for green areas (parks, sport facilities, cemeteries, etc.) in the presence of rabbits, but avoided them when rabbits were absent. General selection of green urban areas is consistent with previous findings of urban habitat selection by European hares in Denmark (Mayer and Sunde 2020), likely because these areas resemble the hares’ preferred habitat, characterized by low vegetation height, providing high-quality forage (Lush, Ward & Wheeler 2017; Mayer *et al*. 2018). Similarly, hares selected for discontinuous urban fabric (often consisting of residential areas) in the absence of rabbits, but avoided them when rabbits co-occurred. Residential gardens, which have been found to constitute important habitats for other urban wildlife (Van Helden *et al*. 2020), might also constitute foraging sites for European hares. It is harder to explain the difference in habitat selection depending on the presence of rabbits that seemingly facilitated the use of green urban areas by European hares (also selected for by rabbits) at the expense of discontinuous urban fabric. One explanation could be that the presence of rabbits increased overall grazing intensity and fertilization via defecation on lawns, leading to increased grass growth, benefitting European hares. This facilitation of European hares by rabbits might be mitigated by dietary differences between the two species (Lush, Ward & Wheeler 2017), allowing their interaction to be rather facilitative than competitive. Similarly, it has been shown that megaherbivore trampling and feeding stimulates high-quality grass regrowth, making it more accessible for smaller ungulates (Wegge, Shrestha & Moe 2006).

We found no evidence that the presence of European hares affected habitat selection by rabbits, indicating that rabbit space use and occurrence was unaffected by hares, as suggested in previous studies (Stott 2003; Katona *et al*. 2004; Flux 2008; Leach, Montgomery & Reid 2017). Rabbits generally selected for green urban areas that likely provided good forage opportunities (Bakker *et al*. 2005). They showed no selection or avoidance for continuous urban fabric, and avoided the other land cover types, including forest. An avoidance of areas that likely provided cover (such as forest and discontinuous urban fabric via hedgerows) might indicate that urban rabbits experienced relaxed predation pressure, as previously proposed, reducing the need for cover (Ziege *et al*. 2016), in combination with these areas probably providing less forage (Lombardi *et al*. 2003). However, as most observations likely came from active rabbits, our results might not apply to inactive rabbits that might select for areas with more cover, leading to a reduced detection probability (Geldmann *et al*. 2016; also see discussion below).

As all 6 urban areas where >10 mountain hare observations were made were located outside the current distribution of the other two lagomorphs, we could not investigate habitat selection depending on species co-occurrence. Mountain hares selected for discontinuous urban fabric, potentially providing both forage and cover, and avoided green urban areas, industry and forest. The apparent avoidance of forest might be related to observer biases (see below). The avoidance of green areas might be related to the absence of cover, as mountain hares are typically associated with habitats providing cover, typically forest (Flux & Angermann 1990; Thulin 2003).

### Study limitations, future considerations, and conclusions

Citizen science data is susceptible to spatial biases with regards to infrastructure and human population density (Geldmann *et al*. 2016). Consequently, citizen observations might have measured human-lagomorph encounters rather than actual habitat preferences, e.g. shown for canids (Mueller, Drake & Allen 2019). Urban areas, while generally having high levels of infrastructure and human population densities, yielding a high sampling effort overall, might still be prone to varying sampling efforts due to being highly heterogeneous (Dickinson, Zuckerberg & Bonter 2010; Crall *et al*. 2011). For example, it is plausible that citizens rather recorded animal observations in their own gardens and in parks compared to city centers and industrial areas. Moreover, detectability also differs between land cover types, accessibility, and depending on animal activity (Mair & Ruete 2016; Pereira-Ribeiro *et al*. 2019). As most observations likely came from active lagomorphs, our results probably represent occurrence and habitat selection of active individuals and from areas that were easily accessible to observers. However, habitat selection by active and inactive lagomorphs differs (Neumann *et al*. 2012; Mayer *et al*. 2018), implying that we might have underestimated the importance of certain land cover types that are predominantly used by resting individuals (e.g. forest patches). Avoiding such biases in citizen observations will be hard. One potential solution would be to select larger spatial scales, as scaling up generally decreases spatial bias and reduces pseudo-absences (Rondinini *et al*. 2006), and to define species occurrence rather than relative abundance. Finally, species might have been misclassified in some cases, resulting in false-positives (Dickinson, Zuckerberg & Bonter 2010). For example, pet rabbits might have been mistaken for wild rabbits, and hybrids of European hares and mountain hares might have been mistaken for either of these two. GPS tagging individuals would enable us to obtain more detailed information on habitat selection and movements by lagomorphs in urban areas, shedding more light on their adaptations to this novel environment. To quantify urban population densities, transect counts could be used (Mayer & Sunde 2020), potentially conducted by citizen scientists if incentivized correctly, like for example the Great Backyard Bird Count (https://www.birdcount.org/).

Our study contributes to the understanding of species co-occurrence patterns and habitat preferences within urban areas, while highlighting the benefits and challenges of citizen science data. We generally found little evidence for competition between the three lagomorphs, though we cannot exclude that urban mountain hare occurrence is inhibited interspecific competition. Future studies should also investigate how the presence of predators, in this case predominantly red foxes (*Vulpes vulpes*), affects the occurrence and habitat selection of lagomorphs within urban areas. Moreover, it would be of interest to shed more light on the drivers of urban colonization by wildlife, to be able to predict urban species occurrence. Insights into species habitat associations within urban areas and depending on co-occurrence with other species can help in targeting urban management plans, which will be useful to identify suitable habitats for desired species and efficient management of pest species (Gaertner *et al*. 2017; Apfelbeck *et al*. 2020).

## Acknowledgements

We thank Henrik Thurfjell for providing insight into Artportalen reports, and Trine Bilde and Trine Wincentz Jensen for comments on an earlier draft of this manuscript.

## Conflict of interest statement

The authors state no conflict of interest.

## Notes

https://doi.org/10.6084/m9.figshare.19699618.v1

### Competing Interest Statement

The authors have declared no competing interest.

## References

Apfelbeck, B., Snep, R.P., Hauck, T.E., Ferguson, J., Holy, M., Jakoby, C., MacIvor, J.S., Schär, L., Taylor, M. & Weisser, W.W. (2020) Designing wildlife-inclusive cities that support human-animal co-existence. Landscape and urban planning,200, 103817.

Arnold, T.W. (2010) Uninformative parameters and model selection using Akaike’s Information Criterion. The Journal of Wildlife Management,74, 1175–1178.

Artdatabanken, S. (2020) Rödlistade arter i Sverige 2020. SLU, Uppsala.

Bakker, E., Reiffers, R., Olff, H. & Gleichman, J. (2005) Experimental manipulation of predation risk and food quality: effect on grazing behaviour in a central-place foraging herbivore. Oecologia,146, 157–167.

Bar-Massada, A. (2015) Complex relationships between species niches and environmental heterogeneity affect species co-occurrence patterns in modelled and real communities. Proceedings of the Royal Society B: Biological Sciences,282, 20150927.

Barton, K. (2020) Package ‘MuMIn’.

Bates, D., Maechler, M., Bolker, B., Walker, S., Christensen, R.H.B., Singmann, H., Dai, B., Eigen, C. & Rcpp, L. (2015) Package ‘lme4’.

Bozek, C.K., Prange, S. & Gehrt, S.D. (2007) The influence of anthropogenic resources on multi-scale habitat selection by raccoons. Urban Ecosystems,10, 413–425.

Brown, L. (2001) Building an Economy for the Earth. Earth Policy Institute.

Calvete, C., Estrada, R., Angulo, E. & Cabezas-Ruiz, S. (2004) Habitat factors related to wild rabbit conservation in an agricultural landscape. Landscape ecology,19, 531–542.

Calvete, C., Estrada, R., Villafuerte, R., Osácar, J. & Lucientes, J. (2002) Epidemiology of viral haemorrhagic disease and myxomatosis in a free-living population of wild rabbits. Veterinary Record,150, 776–782.

Carrete, M., Lambertucci, S.A., Speziale, K., Ceballos, O., Travaini, A., Delibes, M., Hiraldo, F. & Donázar, J.A. (2010) Winners and losers in human-made habitats: interspecific competition outcomes in two Neotropical vultures. Animal Conservation,13, 390–398.

Chambers, L.K. & Dickman, C.R. (2002) Habitat selection of the long-nosed bandicoot, *Perameles nasuta* (Mammalia, Peramelidae), in a patchy urban environment. Austral Ecology,27, 334–342.

Contesse, P., Hegglin, D., Gloor, S., Bontadina, F. & Deplazes, P. (2004) The diet of urban foxes (Vulpes vulpes) and the availability of anthropogenic food in the city of Zurich, Switzerland. Mammalian Biology-Zeitschrift für Säugetierkunde,69, 81–95.

Crall, A.W., Newman, G.J., Stohlgren, T.J., Holfelder, K.A., Graham, J. & Waller, D.M. (2011) Assessing citizen science data quality: an invasive species case study. Conservation Letters,4, 433–442.

Dickinson, J.L., Zuckerberg, B. & Bonter, D.N. (2010) Citizen science as an ecological research tool: challenges and benefits. Annual Review of Ecology, Evolution, and Systematics,41, 149–172.

Duduś, L., Zalewski, A., Kozioł, O., Jakubiec, Z. & Król, N. (2014) Habitat selection by two predators in an urban area: The stone marten and red fox in Wrocław (SW Poland). Mammalian Biology,79, 71–76.

Estevo, C.A., Nagy-Reis, M.B. & Nichols, J.D. (2017) When habitat matters: Habitat preferences can modulate co-occurrence patterns of similar sympatric species. PLoS One,12, e0179489.

Flux, J.E. (1993) Relative effect of cats, myxomatosis, traditional control, or competitors in removing rabbits from islands. New Zealand Journal of Zoology,20, 13–18.

Flux, J.E. (2008) A review of competition between rabbits (Oryctolagus cuniculus) and hares (Lepus europaeus). Lagomorph biology, 241–249.

Flux, J.E. & Angermann, R. (1990) The hares and jackrabbits. Rabbits, hares and pikas. Status survey and conservation action plan,4, 61–94.

Gaertner, M., Wilson, J.R., Cadotte, M.W., MacIvor, J.S., Zenni, R.D. & Richardson, D.M. (2017) Non-native species in urban environments: patterns, processes, impacts and challenges. pp. 3461–3469.Springer.

Geldmann, J., Heilmann-Clausen, J., Holm, T.E., Levinsky, I., Markussen, B., Olsen, K., Rahbek, C. & Tøttrup, A.P. (2016) What determines spatial bias in citizen science? Exploring four recording schemes with different proficiency requirements. Diversity and Distributions,22, 1139–1149.

Grimm, N.B., Faeth, S.H., Golubiewski, N.E., Redman, C.L., Wu, J., Bai, X. & Briggs, J.M. (2008) Global change and the ecology of cities. Science,319, 756–760.

Grueber, C., Nakagawa, S., Laws, R. & Jamieson, I. (2011) Multimodel inference in ecology and evolution: challenges and solutions. Journal of evolutionary biology,24, 699–711.

Haigh, A. & Lawton, C. (2007) Wild mammals of an Irish urban forest. The Irish Naturalists’ Journal, 395–403.

Hijmans, R.J., van Etten, J., Cheng, J., Mattiuzzi, M., Sumner, M., Greenberg, J.A., Lamigueiro, O.P., Bevan, A., Racine, E.B. & Shortridge, A. (2015) Package ‘raster’. R package.

Jansson, G. & Pehrson, Å. (2007) The recent expansion of the brown hare (Lepus europaeus) in Sweden with possible implications to the mountain hare (L. timidus). European Journal of Wildlife Research,53, 125–130.

Katona, K., Bíró, Z., Hahn, I., Kertész, M. & Altbacker, V. (2004) Competition between European hare and European rabbit in a lowland area, Hungary: a long-term ecological study in the period of rabbit extinction. FOLIA ZOOLOGICA-PRAHA-,53, 255–268.

Kelling, S., Fink, D., La Sorte, F.A., Johnston, A., Bruns, N.E. & Hochachka, W.M. (2015) Taking a ‘Big Data’approach to data quality in a citizen science project. Ambio,44, 601–611.

Kohli, B.A., Terry, R.C. & Rowe, R.J. (2018) A trait-based framework for discerning drivers of species co-occurrence across heterogeneous landscapes. Ecography,41, 1921–1933.

Leach, K., Montgomery, W.I. & Reid, N. (2015) Biogeography, macroecology and species’ traits mediate competitive interactions in the order L agomorpha. Mammal Review,45, 88–102.

Leach, K., Montgomery, W.I. & Reid, N. (2016) Modelling the influence of biotic factors on species distribution patterns. Ecological modelling,337, 96–106.

Leach, K., Montgomery, W.I. & Reid, N. (2017) Characterizing biotic interactions within the Order Lagomorpha using Joint Species Distribution Models at 3 different spatial scales. Journal of Mammalogy,98, 1434–1442.

Lees, A.C. & Bell, D.J. (2008) A conservation paradox for the 21st century: the European wild rabbit Oryctolagus cuniculus, an invasive alien and an endangered native species. Mammal Review,38, 304–320.

Levänen, R., Pohjoismäki, J.L. & Kunnasranta, M. (2019) Home ranges of semi-urban brown hares (Lepus europaeus) and mountain hares (Lepus timidus) at northern latitudes. Annales Zoologici Fennici, pp. 107–120.BioOne.

Lombardi, L., Fernández, N., Moreno, S. & Villafuerte, R. (2003) Habitat-related differences in rabbit (Oryctolagus cuniculus) abundance, distribution, and activity. Journal of Mammalogy,84, 26–36.

Luniak, M. (2004) Synurbization–adaptation of animal wildlife to urban development. Proc. 4th Int. Symposium Urban Wildl. Conserv. Tucson, pp. 50–55.Citeseer.

Lush, L., Ward, A. & Wheeler, P. (2017) Dietary niche partitioning between sympatric brown hares and rabbits. Journal of Zoology,303, 36–45.

Magle, S.B., Hunt, V.M., Vernon, M. & Crooks, K.R. (2012) Urban wildlife research: past, present, and future. Biological Conservation,155, 23–32.

Magnusson, A., Skaug, H., Nielsen, A., Berg, C., Kristensen, K., Maechler, M., van Bentham, K., Bolker, B., Brooks, M. & Brooks, M.M. (2017) Package ‘glmmTMB’. R Package Version 0.2. 0.

Mair, L. & Ruete, A. (2016) Explaining spatial variation in the recording effort of citizen science data across multiple taxa. PLoS One,11, e0147796.

Mayer, M. & Sunde, P. (2020) Colonization and habitat selection of a declining farmland species in urban areas. Urban Ecosystems, 23, 543–555.

Mayer, M., Ullmann, W., Sunde, P., Fischer, C. & Blaum, N. (2018) Habitat selection by the European hare in arable landscapes: The importance of small-scale habitat structure for conservation. Ecology and Evolution,8, 11619–11633.

McKinney, M.L. (2006) Urbanization as a major cause of biotic homogenization. Biological Conservation,127, 247–260.

Mueller, M.A., Drake, D. & Allen, M.L. (2019) Using citizen science to inform urban canid management. Landscape and urban planning,189, 362–371.

Møller, A.P. (2012) Urban areas as refuges from predators and flight distance of prey. Behavioral Ecology,23, 1030–1035.

Neumann, F., Schai-Braun, S., Weber, D. & Amrhein, V. (2012) European hares select resting places for providing cover. Hystrix, the Italian Journal of Mammalogy,22.

O’hara, R.B. & Kotze, D.J. (2010) Do not log-transform count data. Methods in Ecology and Evolution,1, 118–122.

Pereira-Ribeiro, J., Ferreguetti, A.C., Bergallo, H.G. & Rocha, C.F.D. (2019) Good timing: evaluating anuran activity and detectability patterns in the Brazilian Atlantic Forest. Wildlife Research,46, 566–572.

Pollock, L.J., Tingley, R., Morris, W.K., Golding, N., O’Hara, R.B., Parris, K.M., Vesk, P.A. & McCarthy, M.A. (2014) Understanding co-occurrence by modelling species simultaneously with a Joint Species Distribution Model (JSDM). Methods in Ecology and Evolution,5, 397–406.

Ramírez-Cruz, G.A., Solano-Zavaleta, I., Mendoza-Hernández, P.E., Méndez-Janovitz, M., Suárez-Rodríguez, M. & Zúñiga-Vega, J.J. (2019) This town ain’t big enough for both of us… or is it? Spatial co-occurrence between exotic and native species in an urban reserve. PLoS One,14, e0211050.

Rondinini, C., Wilson, K.A., Boitani, L., Grantham, H. & Possingham, H.P. (2006) Tradeoffs of different types of species occurrence data for use in systematic conservation planning. Ecology Letters,9, 1136–1145.

Rutz, C. (2008) The establishment of an urban bird population. Journal of Animal Ecology,77, 1008–1019.

Rödel, H.G. & Dekker, J.J. (2012) Influence of weather factors on population dynamics of two lagomorph species based on hunting bag records. European Journal of Wildlife Research,58, 923–932.

Serrano, S. & Hidalgo de Trucios, S. (2011) Burrow types of the European wild rabbit in southwestern Spain. Ethology Ecology & Evolution,23, 81–90.

Shochat, E., Warren, P.S., Faeth, S.H., McIntyre, N.E. & Hope, D. (2006) From patterns to emerging processes in mechanistic urban ecology. Trends in Ecology & Evolution,21, 186–191.

Smith, R.K., Jennings, N.V. & Harris, S. (2005) A quantitative analysis of the abundance and demography of European hares *Lepus europaeus* in relation to habitat type, intensity of agriculture and climate. Mammal Review,35, 1–24.

Stott, P. (2003) Use of space by sympatric European hares (Lepus europaeus) and European rabbits (Oryctolagus cuniculus) in Australia. Mammalian Biology,68, 317–327.

Tapia, L., Domínguez, J., Regos, A. & Vidal, M. (2014) Using remote sensing data to model European wild rabbit (Oryctolagus cuniculus) occurrence in a highly fragmented landscape in northwestern Spain. Acta Theriologica,59, 289–298.

Thulin, C.G. (2003) The distribution of mountain hares Lepus timidus in Europe: a challenge from brown hares L. europaeus? Mammal Review,33, 29–42.

Ulrich, W., Banks-Leite, C., De Coster, G., Habel, J.C., Matheve, H., Newmark, W.D., Tobias, J.A. & Lens, L. (2018) Environmentally and behaviourally mediated co-occurrence of functional traits in bird communities of tropical forest fragments. Oikos,127, 274–284.

Van Helden, B.E., Close, P.G., Stewart, B.A., Speldewinde, P.C. & Comer, S.J. (2020) An underrated habitat: Residential gardens support similar mammal assemblages to urban remnant vegetation. Biological Conservation,250, 108760.

Wagenmakers, E.-J. & Farrell, S. (2004) AIC model selection using Akaike weights. Psychonomic bulletin & review,11, 192–196.

Wegge, P., Shrestha, A.K. & Moe, S.R. (2006) Dry season diets of sympatric ungulates in lowland Nepal: competition and facilitation in alluvial tall grasslands. Ecological research,21, 698–706.

Ziege, M., Babitsch, D., Brix, M., Kriesten, S., Straskraba, S., Wenninger, S., Wronski, T. & Plath, M. (2016) Extended diurnal activity patterns of European rabbits along a rural-to-urban gradient. Mammalian Biology,81, 534–541.

Ziege, M., Brix, M., Schulze, M., Seidemann, A., Straskraba, S., Wenninger, S., Streit, B., Wronski, T. & Plath, M. (2015) From multifamily residences to studio apartments: shifts in burrow structures of E uropean rabbits along a rural-to-urban gradient. Journal of Zoology,295, 286–293.

Ziege, M., Hermann, B.T., Kriesten, S., Merker, S., Ullmann, W., Streit, B., Wenninger, S. & Plath, M. (2020) Ranging behavior of European rabbits (Oryctolagus cuniculus) in urban and suburban landscapes. Mammal Research,65, 607–614.

Zuur, A.F., Ieno, E.N. & Elphick, C.S. (2010) A protocol for data exploration to avoid common statistical problems. Methods in Ecology and Evolution,1, 3–14.

