## Supplementary material for "Co-occurrence patterns and habitat selection of the mountain hare, European hare, and European rabbit in urban areas of Sweden": Table S1

**Table S1:** Table of urban land cover categories used in this study and their corresponding CORINE land cover classes.

| Urban land cover category | CORINE land cover classes |
| --- | --- |
| 1 Continuous urban fabric | 111 Continuous urban fabric |
| 2 Discontinuous urban fabric | 112 Discontinuous urban fabric |
| 3 Industry | 121 Industrial or commercial units 122 Road and rail networks 123 Port areas 124 Airports |
| 4 Green urban areas | 141 Green urban areas |
|  | 142 Sport and leisure facilities |
| 5 Agriculture | 211 Non-irrigated arable land |
|  | 231 Pastures |
|  | 242 Complex cultivation patterns |
|  | 243 Land principally occupied by agriculture, with significant areas of natural vegetation |
| 6 Forest and Heathland | 311 Broad-leaved forest |
|  | 312 Coniferous forest |
|  | 313 Mixed forest |
| 7 Water | 511 Water courses |
|  | 512 Water bodies |
|  | 521 Coastal lagoons |
|  | 522 Estuaries |
|  | 523 Sea and ocean |
| 7 Other areas | 131 Mineral extraction sites 132 Dump sites 133 Construction sites |
|  | 321 Natural grasslands |
|  | 322 Moors and heathland |
|  | 323 Sclerophyllous vegetation |
|  | 324 Transitional woodland-shrub |
|  | 331 Beaches, dunes, sands |
|  | 332 Bare rocks |
|  | 333 Sparsely vegetated areas |
|  | 334 Burnt areas |
|  | 335 Glaciers and perpetual snow |
|  | 411 Inland marshes |
|  | 412 Peat bogs |
|  | 421 Salt marshes |
|  | 422 Salines |
|  | 423 Intertidal flats |
