## Supplementary material for "Co-occurrence patterns and habitat selection of the mountain hare, European hare, and European rabbit in urban areas of Sweden": Table S2

**Table S2:** Overview of the candidate models based on biological hypothesis and the best model for the analysis of number of urban observations, separately for European hares, European rabbits, and mountain hares.

| Analysis/Model |  |  |  |  |  |  |
| --- | --- | --- | --- | --- | --- | --- |
| *European hare observations* | Parameters | df | logLik | AICc | delta AIC | AIC weight |
| Best | Proportion forest + Proportion continuous urban + Proportion surrounding agriculture + Temperature + Precipitation + Size of urban area + Rabbit presence | 9 | -270 | 560 | 0 | 0.99 |
| Climate and size of urban area | Altitude + Temperature + Precipitation + Size of urban area | 6 | -278 | 569 | 9 | 0.01 |
| Co-occurrence of other leporids | Mountain hare presence + Rabbit presence | 4 | -291 | 590 | 30 | 0 |
| Urban land cover | Proportion agriculture + Proportion forest + Proportion continuous urban + Proportion discontinuous urban + Proportion green urban + Proportion industry | 8 | -294 | 606 | 46 | 0 |
| Surrounding land cover | Proportion agriculture + Proportion urban | 4 | -302 | 613 | 53 | 0 |
| *Mountain hare observations* | Parameters | df | logLik | AICc | delta AIC | AIC weight |
| Best | Proportion agriculture + Proportion discontinuous urban + Proportion surrounding urban areas + Altitude + Temperature + Size of urban area + European hare presence | 8 | -125 | 267 | 0 | 0.913 |
| Climate and size of urban area | Altitude + Temperature + Precipitation + Size of urban area | 6 | -129 | 272 | 5 | 0.087 |
| Co-occurrence of other leporids | European hare presence + Rabbit presence | 4 | -167 | 343 | 76 | 0 |
| Urban land cover | Proportion agriculture + Proportion forest + Proportion continuous urban + Proportion discontinuous urban + Proportion green urban + Proportion industry | 4 | -168 | 344 | 77 | 0 |
| Surrounding land cover | Proportion forest + Proportion urban | 8 | -166 | 350 | 83 | 0 |
| *Rabbit observations* | Parameters | df | logLik | AICc | delta AIC | AIC weight |
| Best | Proportion forest + Proportion discontinuous urban + Proportion green urban + Proportion industry + Proportion surrounding urban areas + Altitude + Sand content + Size of urban area | 10 | -176 | 376 | 0 | 0.998 |
| Climate and size of urban area | Altitude + Temperature + Precipitation + Size of urban area + Sand content | 6 | -187 | 388 | 12 | 0.002 |
| Co-occurrence of other leporids | European hare presence + Mountain hare presence | 4 | -192 | 392 | 16 | 0 |
| Urban land cover | Proportion agriculture + Proportion forest + Proportion continuous urban + Proportion discontinuous urban + Proportion green urban + Proportion industry | 8 | -188 | 394 | 18 | 0 |
| Surrounding land cover | Proportion agriculture + Proportion urban | 4 | -194 | 396 | 21 | 0 |
