## Supplementary material for "Co-occurrence patterns and habitat selection of the mountain hare, European hare, and European rabbit in urban areas of Sweden": Table S3

**Table S3:** Estimate, standard error (SE), lower 95% confidence interval (LCI) and upper 95% confidence interval (UCI) of explanatory variables for the analyses of the number of urban observations separately for European hares, mountain hares, and European rabbits. Informative parameters are in bold.

| Parameter | Estimate | SE | LCI | UCI |
| --- | --- | --- | --- | --- |
| *European hares* |  |  |  |  |
| Intercept | 2.48 | 0.15 | 2.18 | 2.77 |
| **Proportion of forest** | **0.48** | **0.15** | **0.19** | **0.77** |
| **Proportion of continuous urban fabric** | **0.34** | **0.18** | **0.00** | **0.69** |
| **Proportion of surrounding agriculture** | **0.71** | **0.21** | **0.29** | **1.12** |
| **Temperature of the coldest quarter** | **-0.44** | **0.22** | **-0.87** | **-0.02** |
| **Precipitation** | **-0.70** | **0.17** | **-1.03** | **-0.38** |
| **Size of the urban area** | **1.29** | **0.25** | **0.79** | **1.78** |
| **Rabbit presence** | **0.43** | **0.18** | **0.07** | **0.79** |
| *Mountain hares* |  |  |  |  |
| Intercept | -0.68 | 0.25 | -1.17 | -0.19 |
| Proportion of agriculture | 0.24 | 0.15 | -0.05 | 0.53 |
| **Proportion of discontinuous urban fabric** | **-0.46** | **0.23** | **-0.91** | **-0.01** |
| **Proportion of surrounding urban areas** | **0.49** | **0.25** | **0.01** | **0.97** |
| **Altitude** | **0.39** | **0.16** | **0.08** | **0.70** |
| **Temperature of the coldest quarter** | **-1.84** | **0.26** | **-2.35** | **-1.32** |
| **Size of the urban area** | **0.66** | **0.22** | **0.23** | **1.08** |
| **Hare presence** | **0.60** | **0.28** | **0.05** | **1.15** |
| *European rabbits* |  |  |  |  |
| Intercept | 1.42 | 0.22 | 1.00 | 1.84 |
| **Proportion of forest** | **-0.61** | **0.27** | **-1.13** | **-0.08** |
| **Proportion of discontinuous urban fabric** | **2.01** | **0.55** | **0.93** | **3.08** |
| **Proportion of green urban areas** | **1.33** | **0.38** | **0.60** | **2.07** |
| **Proportion of industry** | **0.93** | **0.44** | **0.07** | **1.78** |
| **Proportion of surrounding urban areas** | **0.74** | **0.27** | **0.20** | **1.28** |
| **Altitude** | **-0.71** | **0.33** | **-1.36** | **-0.06** |
| **Soil sand content** | **0.75** | **0.39** | **0.00** | **1.51** |
| **Size of the urban area** | **0.82** | **0.28** | **0.26** | **1.37** |
