## Supplementary figures and images for "Co-occurrence patterns and habitat selection of the mountain hare, European hare, and European rabbit in urban areas of Sweden"

### Fig. S1

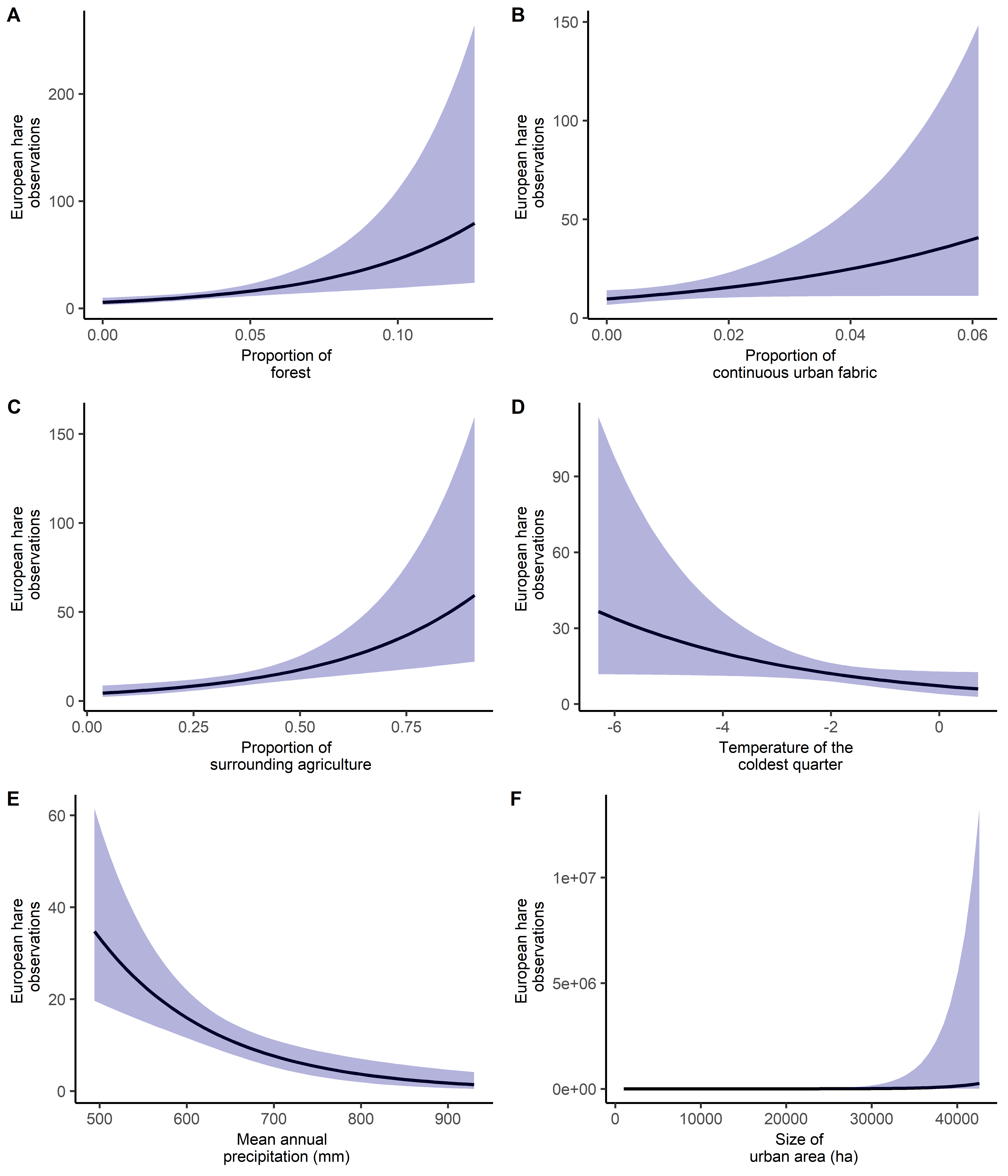

### Fig. S2

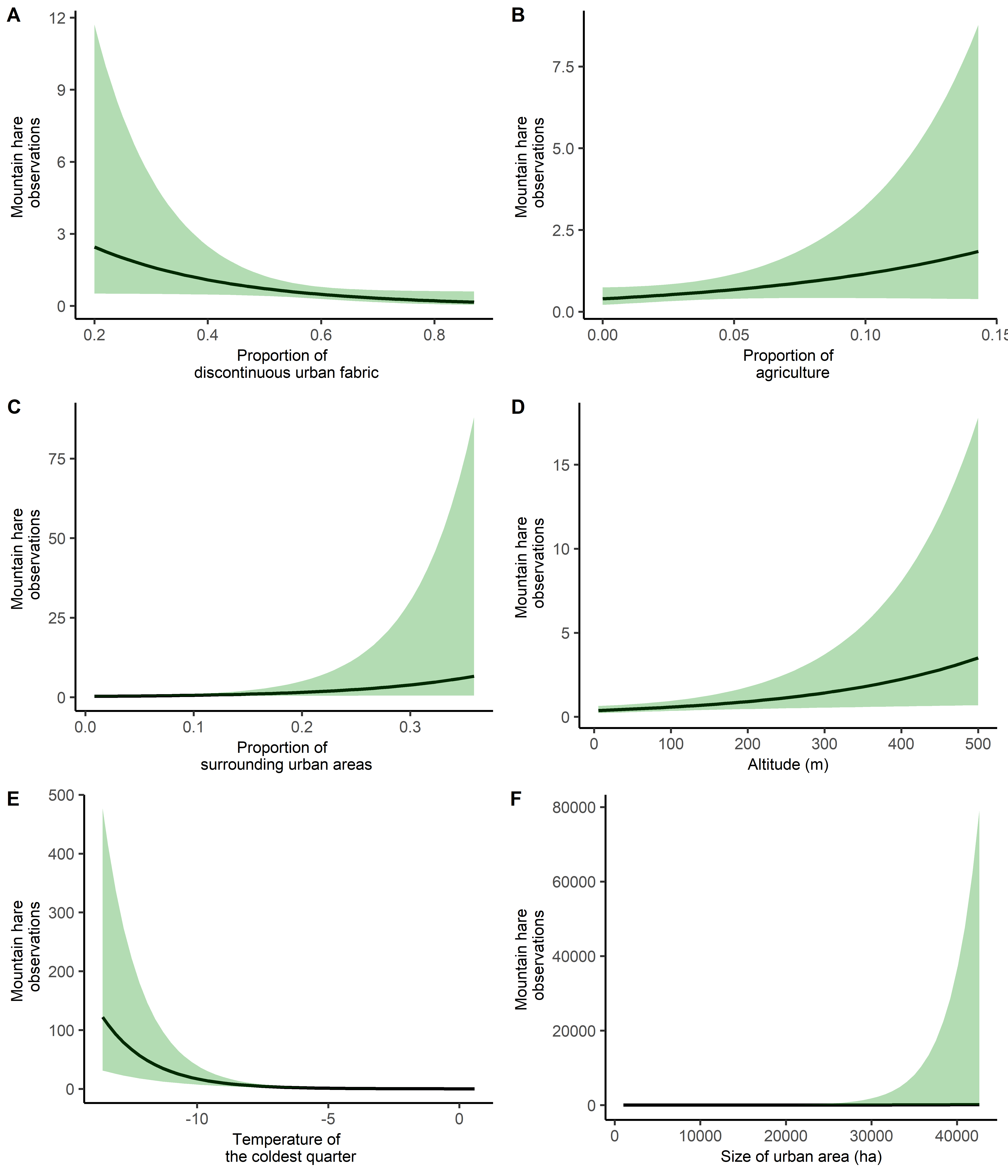

### Fig. S3

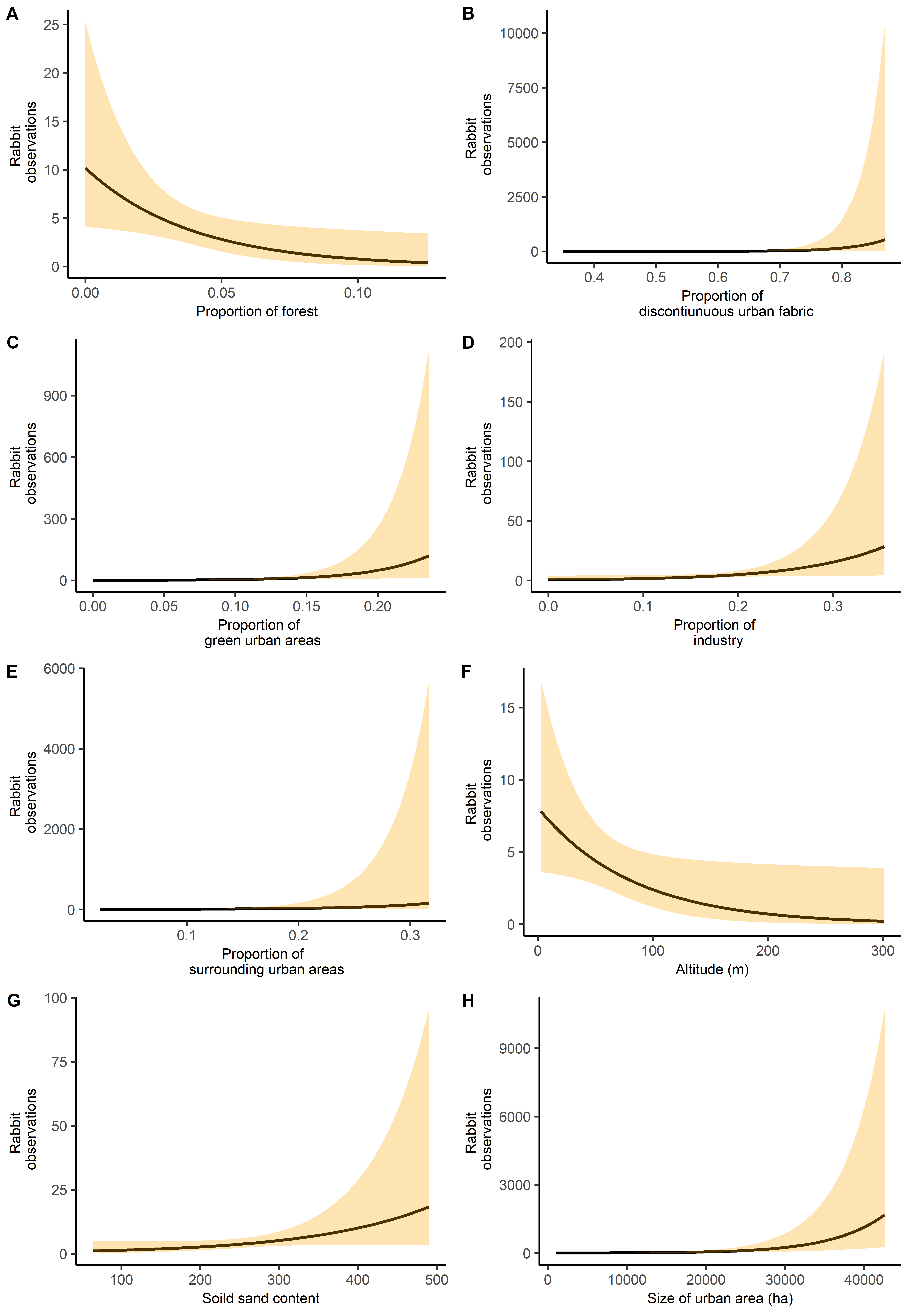
